# Nanopore detection of bacterial DNA base modifications

**DOI:** 10.1101/127100

**Authors:** Alexa B.R. McIntyre, Noah Alexander, Aaron S. Burton, Sarah Castro-Wallace, Charles Y. Chiu, Kristen K. John, Sarah E. Stahl, Sheng Li, Christopher E. Mason

## Abstract

The common bacterial base modification N6-methyladenine (m^6^A) is involved in many pathways related to an organism’s ability to survive and interact with its environment. Recent research has shown that nanopore sequencing can detect m^5^C with per-read accuracy of upwards of 80% but m^6^A with significantly lower accuracy. Here we use a binary classifier to improve m^6^A classification by marking adenines as methylated or unmethylated based on differences between measured and expected current values as each adenine travels through the nanopore. We also illustrate the importance of read quality for base modification detection and compare to PacBio methylation calls. With recent demonstrations of nanopore sequencing in Antarctica and onboard the International Space Station, the ability to reliably characterize m^6^A presents an opportunity to further examine the role of methylation in bacterial adaptation to extreme or very remote environments.

## Introduction

*N*6-methyladenine is the most common modified base in bacterial DNA and plays roles in restriction-modification systems (1), new strand DNA repair (2), and regulation of gene expression (3,4). Methylase mutations and overexpression can affect virulence (5,6) and interactions between bacteria and host cells (7). Target sequences vary by methylase, with the prominent Dam family primarily targeting GATC motifs in *Gammaproteobacteria* and the CcrM family targeting GANTC motifs in *Alphaproteobacteria* (8). In *Escherichia coli*, the adaptive response protein AlkB can demethylate m^6^A sites (9), suggesting dynamic regulation of the mark. The same AlkB machinery also responds to DNA damage, including marks associated with radiation (9,10), although the interaction between damage repair and m^6^A modulation is unknown. Competitive protein binding at methylase target sites can also heritably disrupt methylation (4).

We have recently confirmed that DNA sequencing in space is possible using a nanopore sequencer (11,12). In addition to genomic information, multiple groups have found that distinct, small changes to electrical currents as DNA travels through nanopores can reveal the presence of base modifications (13–15). Methods of detection have so far focused on common mammalian modifications, specifically looking at the eukaryotic contexts of homo-methylated CG-repeats (13) and m^5^C versus m^5^hC (14), but have found lower accuracy (∼70%) for other modifications, including m^6^A (14,15).

Existing methods to detect m^6^A in DNA include sequencing after immunoprecipitation or restriction digests. Immunoprecipitation can reveal methylated areas, but not identify individual bases, while restriction enzyme-based methods are limited to particular motifs and many are unable to differentiate hemimethylation and full methylation (16). PacBio sequencing provides the most precise localization of m^6^A through changes to DNA polymerase kinetics surrounding methylated bases, measured as an inter-pulse duration (6,17–19). Using orthogonal PacBio data generated for samples recently sequenced DNA on the International Space Station (ISS), we identified sets of > 90% methylated and 0% methylated positions in the *E. coli* MG1655 K12 genome to train and test binary classifiers to detect m^6^A in nanopore data. Here, we present a new nucleic acid modification caller (mCaller) for the detection of m^6^A in single reads from deviations between measured and expected signals, now available at github.com/al-mcintyre/mcaller.

## Results

We first demonstrate that the effects of DNA methylation on template strand sequence quality vary depending on the version of the technology and base caller used (Supplementary Figure 1). Oxford Nanopore Technologies provides an estimated mean and standard deviation for currents corresponding to individual 5- or 6-mers, which were used for base calling with hidden Markov models (20) before Oxford Nanopore switched to using recurrent neural networks. These distributions assume that five or six bases in and around the pore affect the current through the pore at a given time point. Previous modification callers built similar distributions for methylated 6-mers, although Rand et al. (2017) noted that these distributions were less distinguishable for m^6^A than m^5^C.

However, across the extended sequence context surrounding an individual methylated site, there are notable shifts from the model that differ in magnitude and direction depending on context (Figure 1A-B). We looked at current deviation (observed - expected values) over sliding windows of six 6-mers around methylated and unmethylated adenines, where each sliding window covers the 11-mer around an adenine. We were initially concerned that longer sequence contexts would require prohibitive amounts of data for training, since there are over a million possible 11-mers (4^10^) that contain only a single central modified position. Yet, some informative patterns are shared among contexts; for R9.4 data, methylation at the 1^st^, 2^nd^, 4^th^, and 5^th^ positions of a 6-mer tends to increase the current with respect to the model values, while methylation at the 3^rd^ or 6^th^ position shows the reverse trend (Figure 1C). We used these current deviations as features to train classifiers.

**Figure 1.**
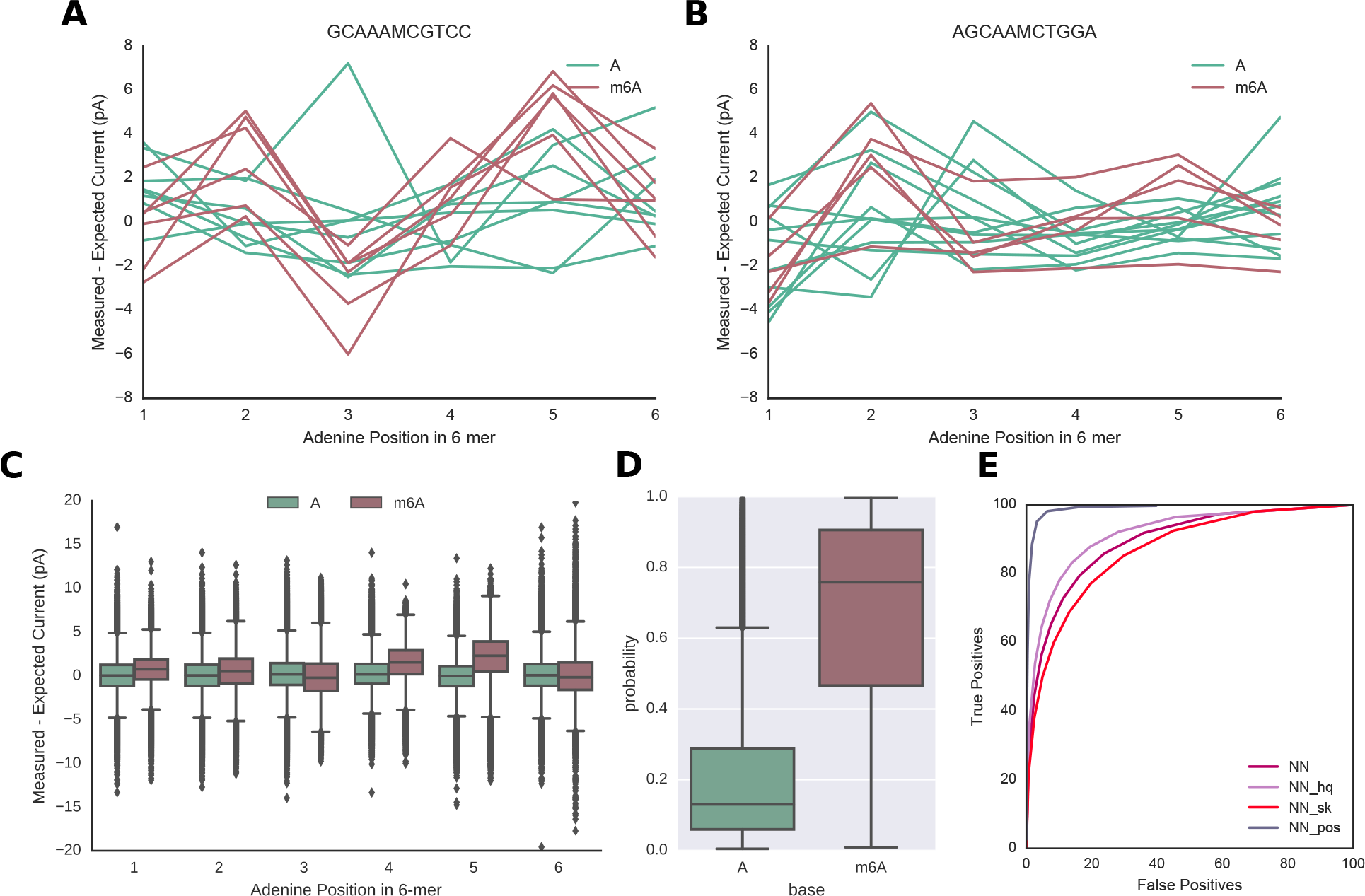
**(A-B)** Changes to current deviations from model values differ depending on the sequence context of a methylated base, with changes more evident in some contexts than others. Shown are two 11-mers for which the central base (marked “M”) was identified as both methylated and unmethylated at different positions in the genome by PacBio. **(C)** General patterns emerge, depending on the position of a methylated base in a 6-mer. For R9.4 data, methylation at the 1^st^, 2^nd^, 4^th^, and 5^th^ positions of a 6-mer tend to increase current measurements with respect to the model, while methylation at the 3^rd^ or 6^th^ position contributes to a slight overall decrease in current. **(D)** Probabilities of methylation defined by a neural network classifier for methylated compared to unmethylated positions in *E. coli*, with a model trained on one dataset and tested on a second. **(E)** ROC curves for the neural network model using R9.4 data, showing true positive rate (methylated positions correctly identified) as a function of false positive rate (unmethylated positions called as methylated) as we vary the probability threshold for classification. We tested modifications to the standard model (“NN”) using only high quality reads (average base quality > 9, “NN_hq”) and classifying observations that included a maximum of two skips (“NN_sk”). A per-position curve (“NN_pos”) was calculated for genomic positions with ≥15X coverage by varying the fraction of reads with probability of methylation scores ≥0.5 required to define a position as methylated.

A neural network classifier produced the highest accuracy among four different methods, including random forest, naïve Bayes, and logistic regression (Supplementary Figure 2), with 81% accuracy using all quality levels of reads (Figure 1D, Supplementary Table 1). The Spearman correlation between the probability estimates from the top two predictors, neural network and random forest, was high, at 0.92 (Supplementary Figure 2D). A receiver operator characteristic curve shows the changes in accuracy at different levels of probability (Figure 1E). Accuracy improved to 84% for higher quality reads and decreased to 78% when we allowed the 11-mers to include a maximum of two skips, or 6-mers for which the sequencer missed recording a current value. When we used a minimum of 15X coverage of the sites, the classifier achieved 94% accuracy and 0.99 AUC. We then tested the hypothesis that methylation would affect a similar range of surrounding current levels as the canonical bases in the Oxford Nanopore models (six). Indeed, taking current differences for four or eight 6-mers surrounding a base reduced classification accuracy (Supplementary Figure 2A,C).

For further validation, we compared methylation predictions at sites called as partially modified (0-90]% through PacBio sequencing, and found a Pearson correlation coefficient of 0.68 between the fractions of methylated bases at each position detected by each method (Figure 2). The nanopore-based method identified less methylation at some sites, however, it is worth noting that PacBio methylation calls in single strands are not as accurate as per-position calls, which could affect estimates of the fraction of reads methylated (18).

**Figure 2.**
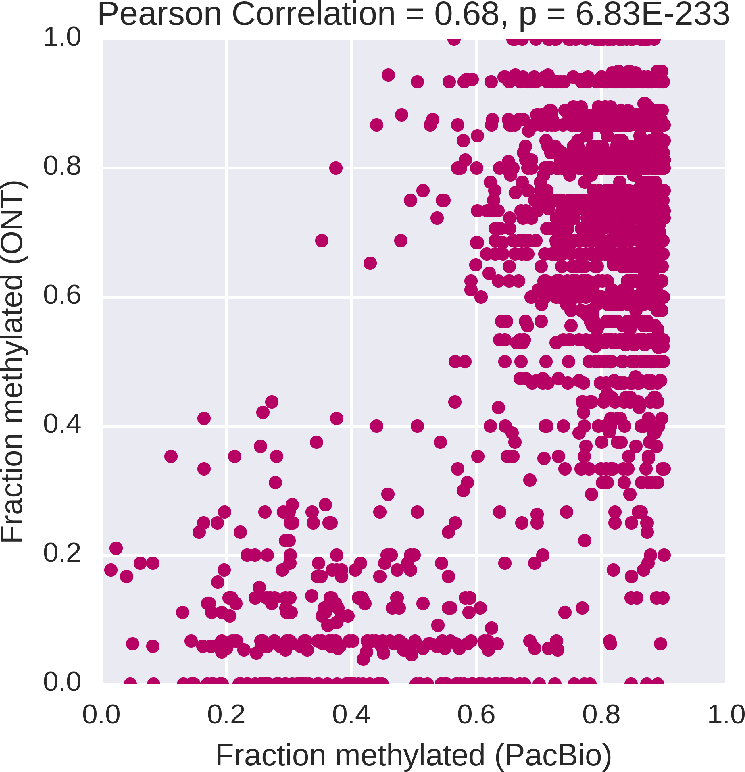
A comparison between the fraction of reads classified as methylated using nanopore data (with a minimum coverage of 15 reads) and PacBio at partially-methylation positions.

For bacterial species like *Escherichia coli* in which one enzyme (Dam) is responsible for most methylation and primarily targets GATC motifs, the similarity in sequence contexts could increase concordance among classification features across sites. Therefore, we also tested the method on a second base modification found in more variable contexts, m^5^C. We used PCR-amplified and M. Sssl-methylated from Simpson et al. (2017) to train and test the method for the identification of 5-methylcytosine in CG contexts. We again found features sufficiently similar across contexts for prediction, with accuracy around 77% (Supplementary Figure 3).

## Methods

Data were generated for nine runs in flight on the ISS, and nine matched control runs on the ground, as described previously (12). The 18 experiments used the same batch of R7.3 flow cells and the same libraries of DNA containing equal masses of native mouse and *E. coli* MG1655 K12 DNA, and m^6^A-free *Lambda phage* DNA. Two additional datasets using the same mixture and newer version R9.4 flow cells were generated at Weill Cornell Medicine (Mason lab) and University of California, San Francisco (Chiu lab).

Orthogonal PacBio data for the same *E. coli* strain were also generated and a single contig was assembled using HGAP v2 (12). Reads were realigned to the *de novo* assembly and methylated positions identified through differences between measured and expected inter-pulse durations using SMRTPortal v2.3.0-RS_Modification_and_Motif_analysis.1 (Supplementary Figure 4) (18,19). All methylated positions included in our analysis had QV scores ≥ 20 (p-values ≤ 0.01) for both the significance of the difference between the inter-pulse duration and the expected background and for the confidence in the modification type (“identificationQv”). We considered positions fully methylated if they had an estimated modification fraction > 90%, and partially methylated if the fraction was > 0 and ≤ 90%. A control set of the same number of positions was chosen at random from adenines not called as partially or fully methylated at any score. No regions of the *E. coli* genome that also align to the *Lambda phage* genome were considered to avoid uncertainty in the provenance of reads.

Nanopore reads were aligned to the genome from PacBio using GraphMap (21). We began development for R7.3 data re-aligning signals to the genome as part of mCaller. First, the signal currents were scaled to the model using scale, drift, and shift parameters, as previously described (22). Next, anchor points were identified on each side of a potentially methylated position at which a read 6-mer and a reference 6-mer match exactly and assumed to have a 100% match probability. A hidden Markov model and the Viterbi algorithm were then used to connect currents to reference 6-mers between the two anchor points. Although this worked well for R7.3 data and helped eliminate noisier reads with no anchor points within a 50 bases of a methylated base, we wanted to test the model using all reads as the technology improved in accuracy, therefore used an existing program, nanopolish eventalign (13,23), to re-align the R9 current data. Besides the figure showing base quality, we focus on the R9.4 data as more indicative of current nanopore performance. Average read quality was added as an additional classification feature to better control for variation among experiments. By default, we did not classify observations with skips.

All binary classification models were tested using scikit-learn implementations (24). Parameters for the classifiers were adjusted using random grid search and 5-fold cross-validation. The highest performing sets of parameters for each algorithm were used to test on a second dataset. ROC plots were generated by successively increasing the probability threshold for identification of methylation, where all positions in the test dataset were classified as either methylated or unmethylated.

## Discussion

For bacteria and other kingdoms of life, single molecule sequencing with PacBio or nanopore can reveal both genomic and epigenomic states of nucleic acids. We show here that the most common bacterial base modification can be detected at 84% accuracy in single reads using picoampere-scale changes in current as individual DNA molecules travel through a nanopore, and at 94% accuracy with 15X or higher coverage. Our strategy has several advantages over previously published work, with accuracy for m^6^A calling in single strands notably improved over the method recently published by Rand, et al. (2017, 70%). Stoiber et al. (2016) required 20X coverage and both native DNA and whole-genome amplification of the same sample to approximately localize base modifications, while our method requires only a model trained to detect the base modification in question to precisely call sites in single reads. Our program is the first designed for training using orthogonally validated positions, rather than artificially methylated or unmethylated reads. It is also context-naïve in that it does not rely on the observation of a particular k-mer context within the training dataset to make a prediction.

Nanopore sequencing is far more portable than traditional sequencing methods. Recent use during the Ebola outbreak in West Africa (25,26) and the zika outbreak in South America (27,28), as well as in the most distant of human-inhabited locales beyond Earth (12), demonstrates new applications for rapid and remote sequencing and diagnostics. As the duration and distance of spaceflight grows, measures are necessary to protect crewmembers from infection. Extensive research describes species-dependent changes to bacterial growth, gene expression, and pathogenicity in microgravity (29–33), in addition to potentially higher mutation rates in space (34). With concurrent impairment of host immune responses (29) and in the confined quarters of a vessel (35), bacteria present an increased hazard. How bacterial genomes change over many generations in space and whether epigenomic changes relate to shifts in gene expression and metabolite production remain unclear, although the latter has been observed in response to other environmental stimuli (36). The portability of nanopore sequencers can enable further research and potential diagnostics, regardless of where such variation occurs.

## Supplementary Information

**Supplementary Figure 1.**
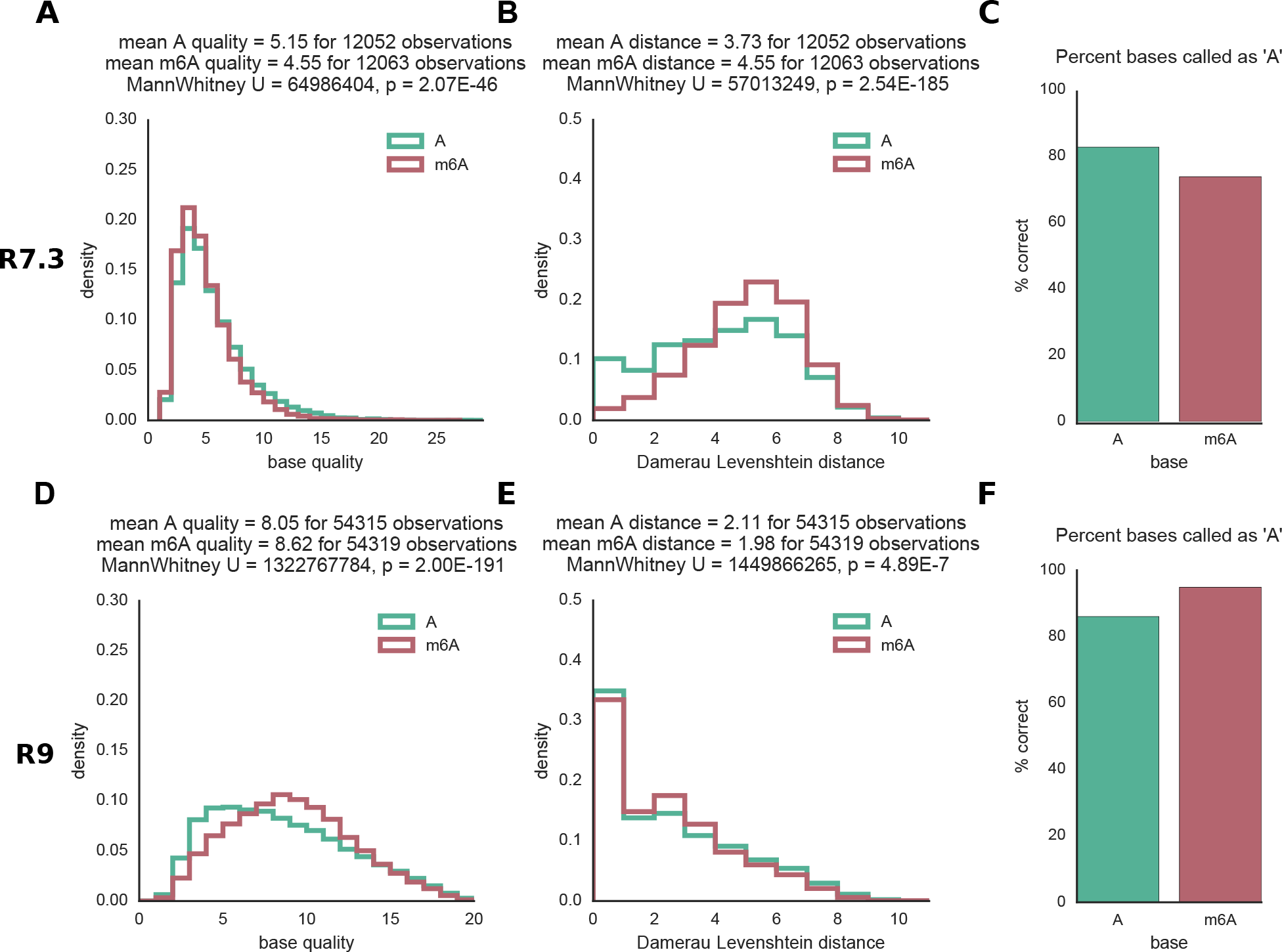
(**A,D**) Methylation at a position has little effect on base quality compared to unmethylated controls (randomly selected from adenines between 20 and 100 bases from a methylated position within the same read). (**B,E**) The edit distance between a base called sequence surrounding a methylated position and a reference genome increases for R7.3 data, but decreases for R9.4 data. (**C,F**) The percentage of bases at methylated vs. unmethylated positions correctly classified as adenine. (**A-C**) R7.3 data from the 2016-08-26 flight, basecalled using a hidden Markov model from Metrichor, (**D-F**) R9.4 data from the Mason lab, basecalled using a recurrent neural network from Metrichor.

**Supplementary Figure 2.**
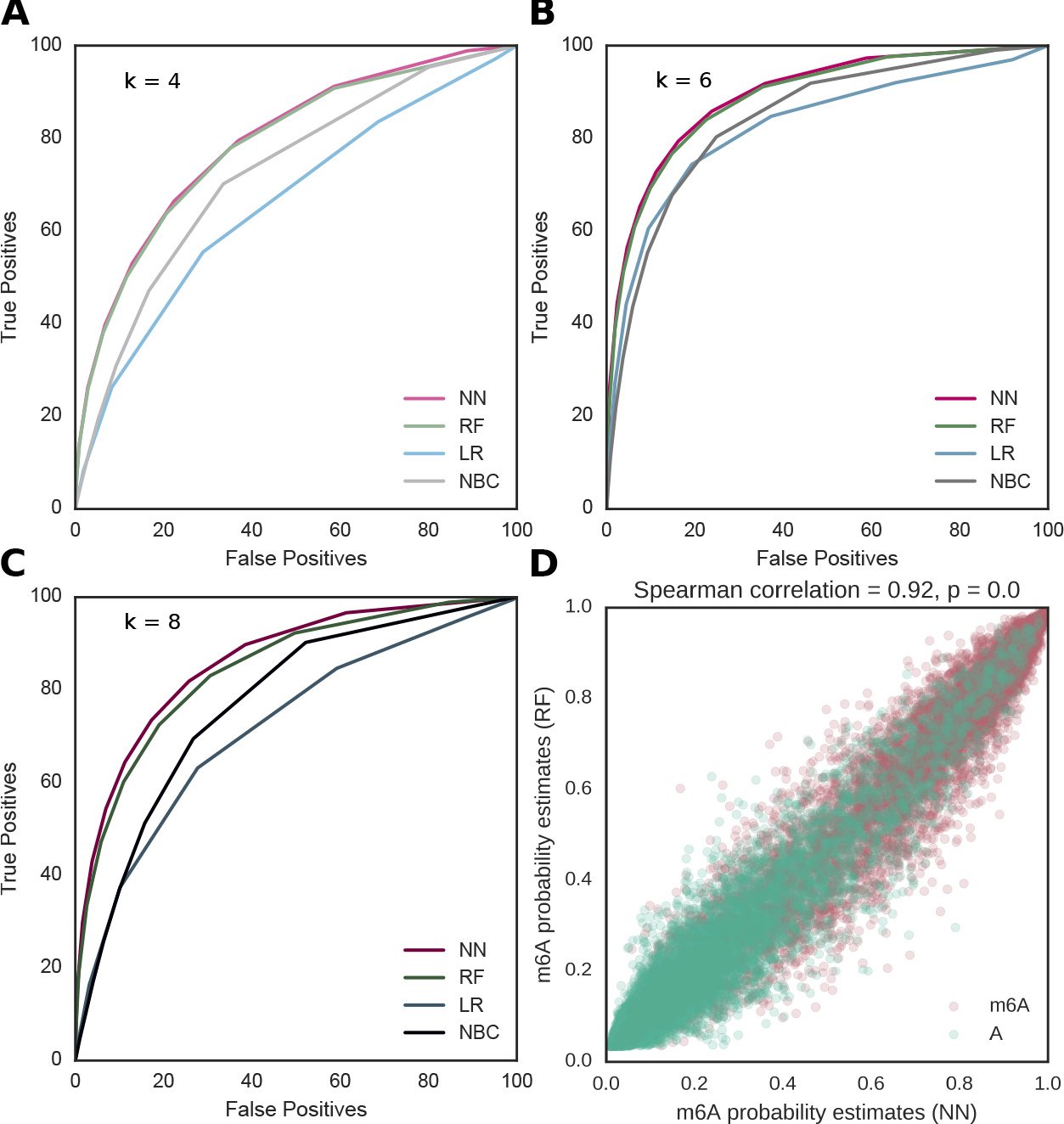
ROC plots for a support multi-layer perceptron neural network (NN), random forest classifier (RF), linear regression classifier (LR), and naïve Bayes classifier (NBC) using (**A**) 7-mer, (**B**) 11-mer, and (**C**) 15-mer contexts to classify. (**D**) A comparison of the probability scores for classifications using the top (x-axis, neural network) and second place (x-axis, random forest classifier) algorithms for 11-mer contexts. Models were generated using one R9.4 experiment and tested on data from a second.

**Supplementary Table 1.** Accuracy and area under the curve (AUC) for methods and parameters tested (number of features and filters) in Figure 1D and Supplementary Figure 2. Methods include neural network (NN), random forest (RF), linear regression (LR), and naïve Bayes classifer (NBC), and filters allowing up to two skips (sk), only high-quality reads (hq) of QV>9, and positions (pos) covered by 15 or more reads.

| # Features | Classifier | Accuracy | AUC |
| --- | --- | --- | --- |
| 4 | NN | 70.1 | 0.798 |
| 4 | RF | 69.2 | 0.793 |
| 4 | LR | 59 | 0.67 |
| 4 | NBC | 60.8 | 0.738 |
| 6 | NN | 80.7 | 0.897 |
| 6 | RF | 79.4 | 0.891 |
| 6 | LR | 75.5 | 0.83 |
| 6 | NBC | 73 | 0.849 |
| 8 | NN | 76.6 | 0.866 |
| 8 | RF | 74.6 | 0.849 |
| 8 | LR | 63.6 | 0.728 |
| 8 | NBC | 63.5 | 0.782 |
| 6 | NN_sk | 77.8 | 0.87 |
| 6 | NN_hq_sk | 80.5 | 0.898 |
| 6 | NN_hq | 84 | 0.921 |
| 6 | NN_pos | 93.4 | 0.989 |

**Supplementary Figure 3.**
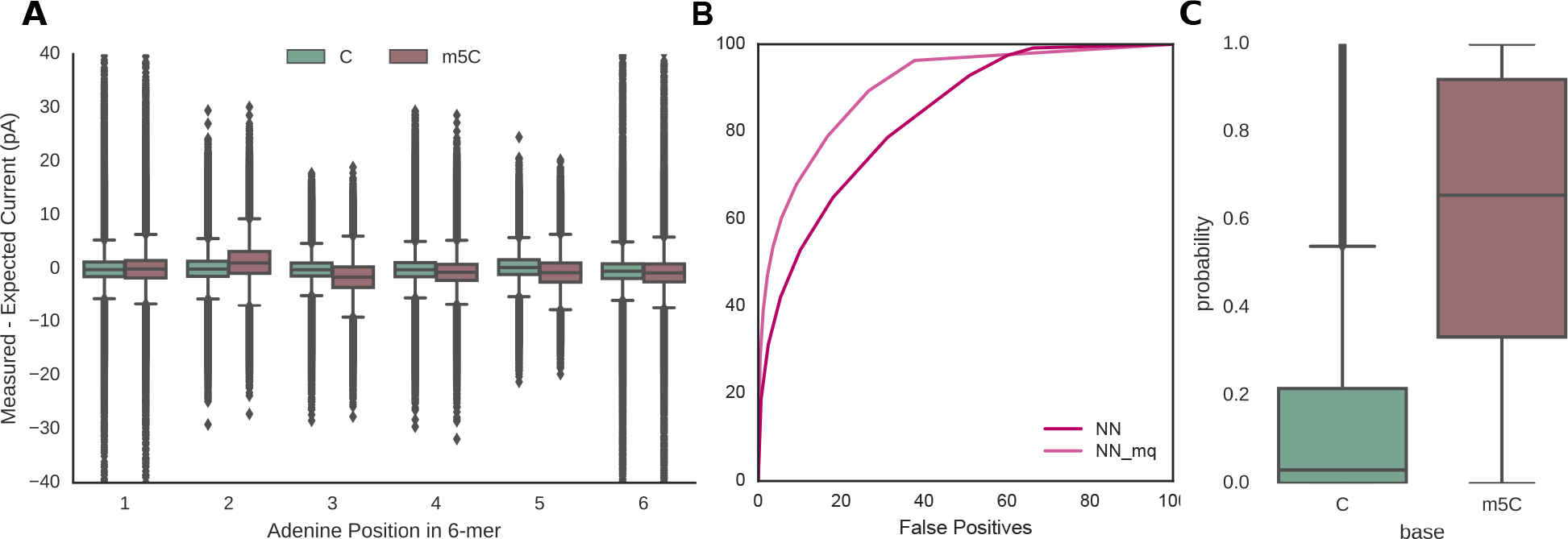
**(A)** Differences between measured and expected current values across different positions in a 6-mer, depending on cytosine methylation status. The variances for both methylated and unmethylated cytosines are higher than for m^6^A, likely in part because the reads were of lower average quality (5.64 for the M.Sssl dataset and 6.41 for the PCR dataset, vs. 7.65 for the Chiu dataset and 7.32 for the Mason dataset). **(B)** A ROC plot shows lower accuracy for m^5^C classifications (73%) than m^6^A. However, limiting the data to “medium quality” (“mq”, > mean base quality score of 7) improves classification to 77%, similar to m^6^A despite including more sites with multiple methylated bases. **(C)** Probability scores for the detection of m^5^C using the same random forest classifier trained and tested using medium quality reads from M. Sssl-modified data and whole genome amplified data.

**Supplementary Figure 4.**
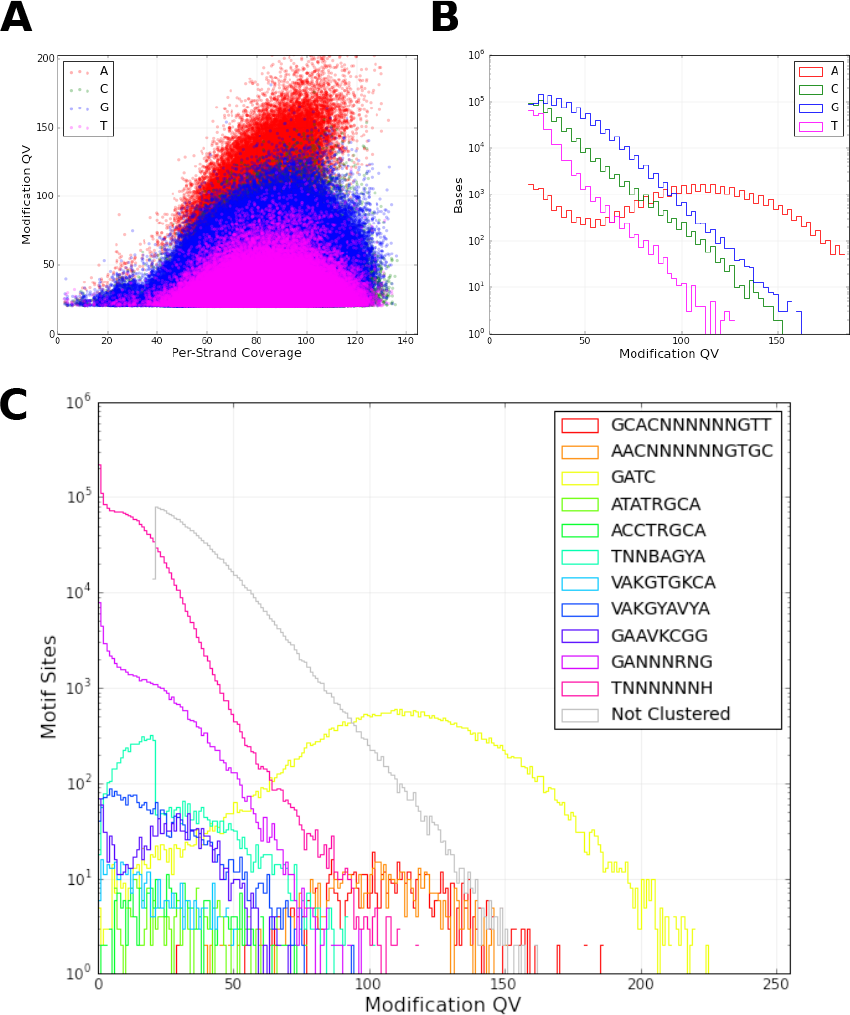
The base modification report from PacBio. **(A)** Modification QV scores for different levels of coverage show adenines form a higher confidence cluster. **(B)** A histogram again shows a higher peak at higher modification QV scores for adenine. **(C)** The motif analysis identified methylation within the recognition sites for Dam (GATC) and the MEcoK methylase (AAC(N_6_)GTGC and GCAC(N_6_)GTT).

